# From prevalence to incidence - a new approach in the hospital setting

**DOI:** 10.1101/554725

**Authors:** Niklas Willrich, Sebastian Haller, Tim Eckmanns, Benedikt Zacher, Tommi Kärki, Diamantis Plachouras, Alessandro Cassini, Carl Suetens, Michael Behnke, Petra Gastmeier, Jan Walter

## Abstract

Point-prevalence surveys (PPSs) are often used to estimate the prevalence of healthcare-associated infections (HAIs). Methods for estimating incidence of HAIs from prevalence have been developed, but application of these methods is often difficult because key quantities, like the average length of infection, cannot be derived directly from the data available in a PPS. We propose a new theory-based method to estimate incidence from prevalence data dealing with these limitations and compare it to other estimation methods in a simulation study. In contrast to previous methods, our method does not depend on any assumptions on the underlying distributions of length of infection and length of stay. As a basis for the simulation study we use data from the second study of nosocomial infections in Germany (Nosokomiale Infektionen in Deutschland, Erfassung und Prävention - NIDEP2) and the European surveillance of HAIs in intensive care units (HAI-Net ICU). The new method compares favourably with the other estimation methods and has the advantage of being consistent in its behaviour across the different setups. It is implemented in an R-package prevtoinc which will be freely available on CRAN (http://cran.r-project.org/).

## INTRODUCTION

Epidemiological information on healthcare-associated infections (HAIs) is often acquired by means of point-prevalence surveys (PPSs). Large-scale PPSs are regularly performed by the European Centre for Disease Prevention and Control (ECDC) (1, 2), as well as the US Centers for Disease Prevention and Control (CDC) (3, 4). While the prevalence of HAIs is an important measure in itself, epidemiologists are usually more interested in the incidence of HAIs. For example, estimations of the burden of HAIs often rely on incidence rather than prevalence (5). Therefore, methods of estimating the incidence rates from the data of PPSs are needed. Under general conditions, the incidence and prevalence can be estimated from one another (6). The question of estimating incidence from prevalence in the context of HAIs has been addressed in the 1980s by two articles (7, 8). The method developed by Rhame and Sudderth (7) is the most commonly applied method for estimating incidence from prevalence (1–3, 5, 9–14). This method however has several limitations:

The Rhame-Sudderth formula was developed using a definition of prevalence that included active and cured infections on the day of the PPS and that is different from the one usually applied in PPSs of HAIs. Another problem with the application of the formula is that it requires a method to estimate the average length of stay and the average length of infection based on data available on the day of the PPS. Without estimates of these quantities from other sources, the application of the estimation method is challenging, because usually only the data obtained on the day of the PPS are available.

In this article, we propose a novel approach dealing with these limitations of estimating incidence from prevalence of HAIs. The proposed approach uses state-of-the-art statistical techniques to estimate the average length of infection and average length of stay in the whole patient population from samples of lengths of infection and hospital stay up to the day of the PPS without relying on any assumption about the distributions of these quantities. We evaluated the new method by comparing it with existing procedures in the literature through simulation studies based on data from the second study of nosocomial infections in Germany (Nosokomiale Infektionen in Deutschland, Erfassung und Prävention - NIDEP2) (15) and from the European surveillance of HAIs in intensive care units (HAI-Net ICU) (16, 17), as well as theoretical distributions.

## METHODS

### Notation

In general, we used the variable *X* to indicate a randomly sampled duration from the whole population and *L* for a randomly sampled duration from the PPS (the duration for a randomly selected patient included in the PPS). *L* is expected to be on average larger than *X*, due to the phenomenon of length-biased sampling (8, 18). We used *A* for the observed duration up to a fixed time for a randomly selected patient at that time point. This was applied to the length of stay and the length of infection.

We used *X, A* and *L* when it was not important to distinguish between the length of stay and the length of infection from a theoretical perspective.

The different concepts for the durations are illustrated in Fig. 1. The notation used in this article is explained in Table 1.

**Fig. 1.**
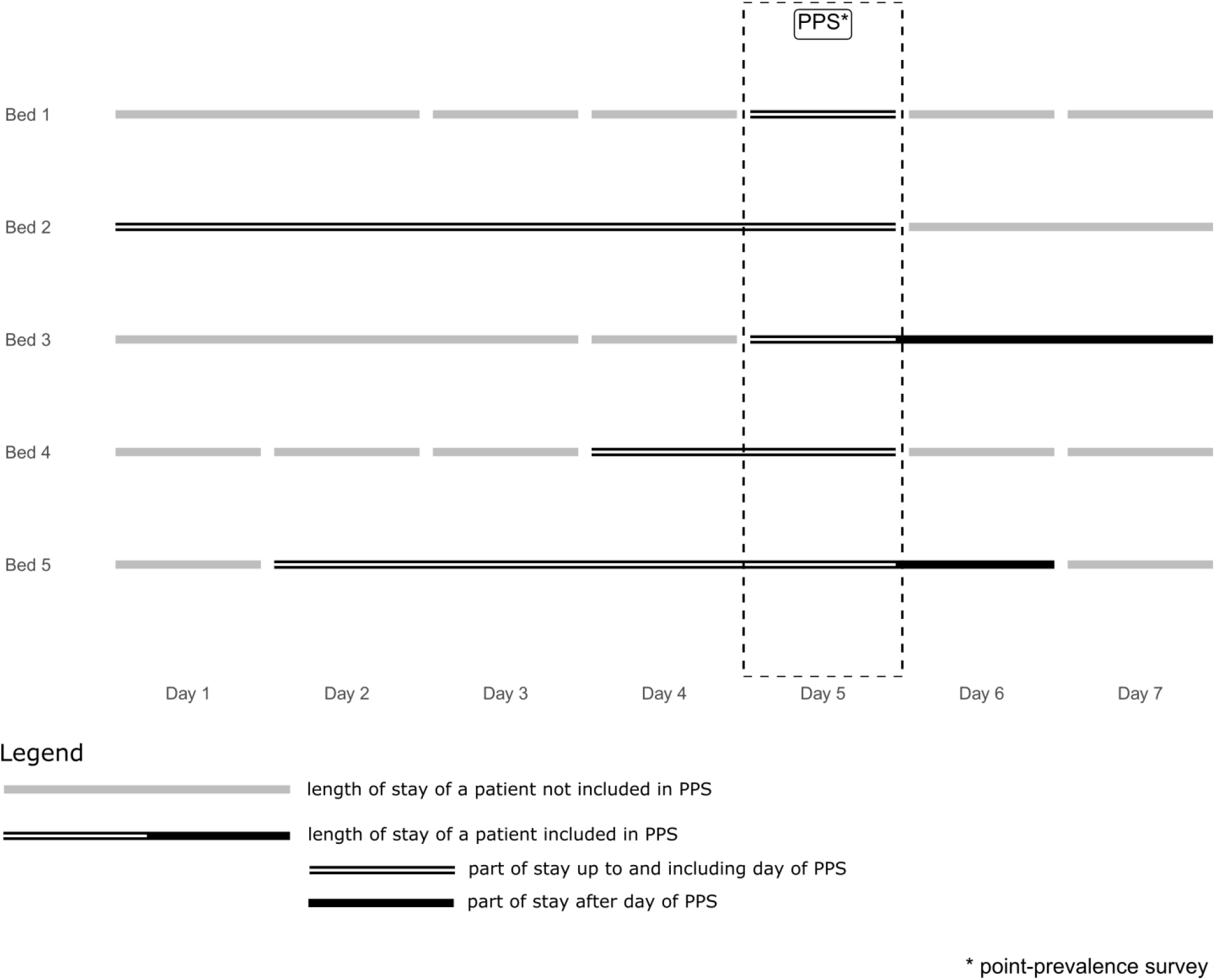
Illustration of different samplings for a hypothetical hospital: Line segments represent patients admitted in a hypothetical hospital. Sampling from X means selecting one of all the line segments at random, sampling from L means only sampling among the segments which intersect with the survey and sampling A means only sampling from the striped parts of these segments representing the part of stay up to and including the day of the PPS.

**Table 1.**
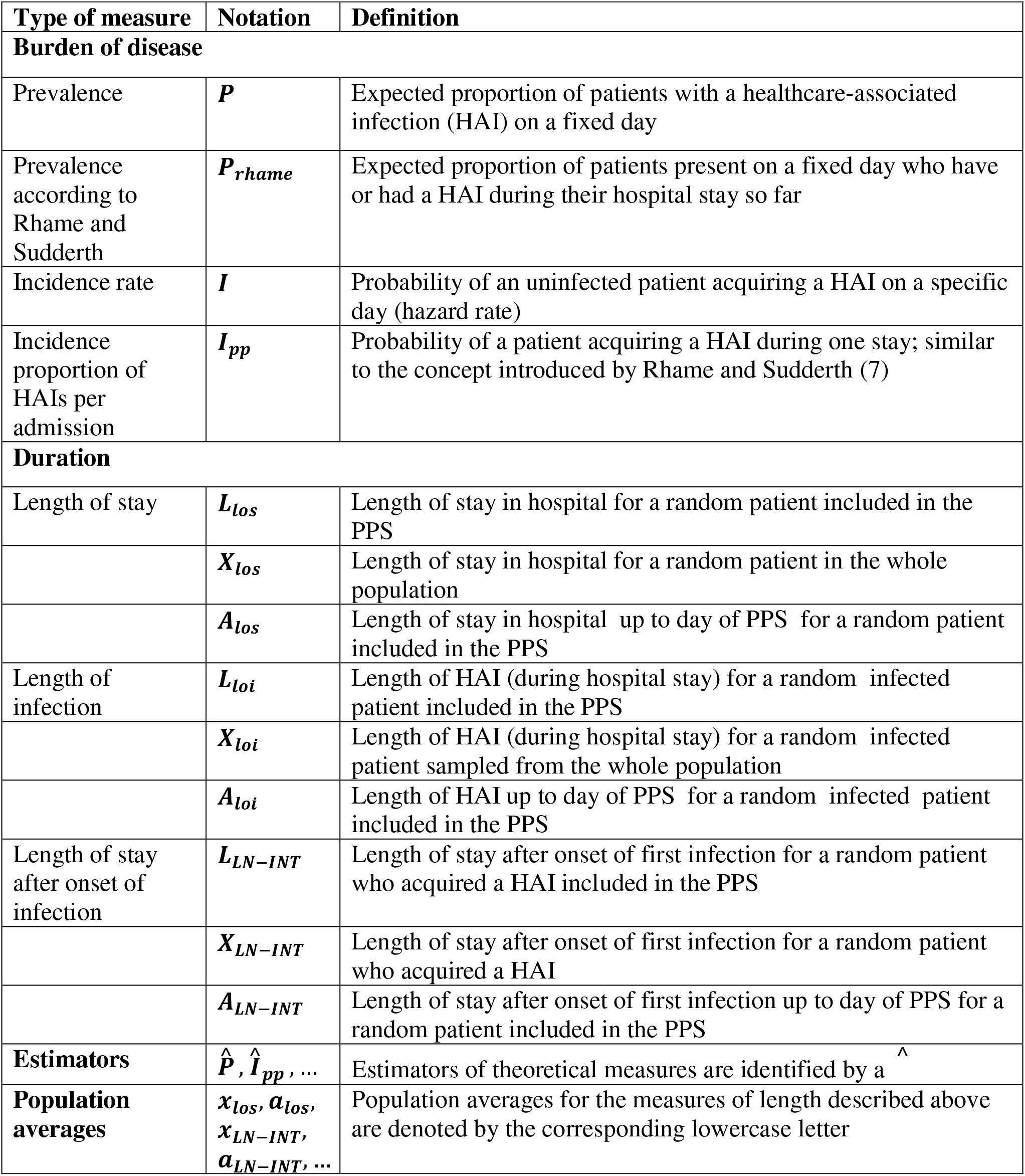
Notation used in this article

### Rhame and Sudderth formula

In line with previous authors (7, 8), we assumed that the patient population is in steady state, i.e. the distribution of characteristics of our sample of patients does not depend on the specific day of the survey.

The original formula of Rhame and Sudderth (7) for the incidence per admission *I_pp_* (slightly simplified and adapted to our notation) is:

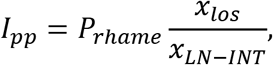

where *x_los_* denotes the average length of stay of a patient, *x_LN-INT_* is the average length of stay for patients after they acquire their first HAI and *I_pp_* is the estimate of the incidence per admission. In this original formulation, *P_rname_* is calculated by counting all patients who had at least one HAI *up to* the time of the survey (and not just the patients that have an active HAI on the date of the survey) and dividing by the total number of patients.

As pointed out above there are two points that complicate the application of this formula in this form:

1. often the PPSs only count patients with *active* infections on the day of the PPS. In these cases, theoretical considerations then require that the term *x_LN-INT_* is replaced by a term *x_loi_* which gives the average length of a HAI (see supplement S1).
2. samples of *X_los_, X_loi_* (or *X_LN-INT_*) are often not available and only the length of stay *A_los_* and possible length of infection *A_loi_* up to the day of the PPS are available.

### New approach

To estimate the distributions of length of stay and length of infection from the observed lengths of stay up to the day of the PPS, we proceeded in two steps:

- We estimated the distributions of length of stay and length of infection up to the day of the PPS (in our notation *A_los_* and *A_loi_*) from the available data,
- We calculated from these distributions the expected lengths (*x_los_* and *x_loi_*) for the whole population.

For the first part, we used an estimator which ensures the monotonicity of the estimated distribution, because the distribution of *A* is always monotonously decreasing. This can be demonstrated by the timeline of occupancy of a hypothetical bed: on average there will always be more patients for whom it is the first day of their stay than the second day, more patients for whom it is the second day of their stay than the third and so on. We use a *Grenander* estimator for discrete distributions described and studied by Jankowski and Wellner (19). This estimator is the maximum likelihood estimator for a discrete monotonously decreasing distribution and is therefore a canonical choice. It is well-studied from a theoretical point and has good properties like consistency and 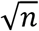 - rate of convergence(19)

Following Freeman and Hutchison (8), in the steady state the relation between prevalence *P* and incidence rate *I* can be written as:

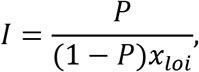

where *x_loi_* is the average length of a HAI in the whole population. To get this equation into the form of Rhame and Sudderth (7), we multiply by the expected length of stay of random patients that are susceptible to an infection (1 − *P*)*x_los_* and get

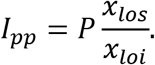

To express *x_loi_* in terms of *A_loi_*, we note the following formula with *N_pat_* the total number of patients at the hospital on the survey day:

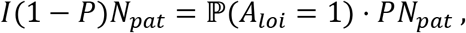

where (1 − *P*)*N_pat_* is the average number of patients at risk, 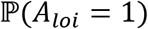 is the average proportion of patients with HAI on the first day of infection and *PN_pat_* is the average number of patients with a HAI.

Both sides of the equation represent the number of average new infections per day; the left hand side as the incidence rate *I* per patient-day-at-risk times the number of patients at risk (1 − *P*)*N_pat_* and the right hand side as the average number of HAI cases on the first day of infection. Therefore

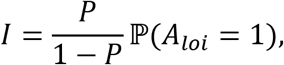

where 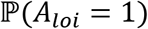 is the probability of sampling *A_loi_* = 1. By comparing with the original incidence rate formula, this gives us, the simple relation

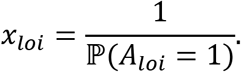

An alternative, more formal route to the formula is based on renewal theory (8) and specifically Eqn. 2.16 from Haviv (20).

This leads to the estimator:

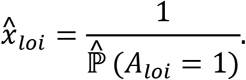

with 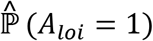 an estimator of 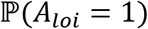. We call 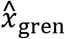 the estimator for *x* based on this procedure with the Grenander estimator (19) for 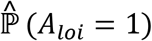.

The general method is equally applicable for the estimation of *x_los_, x_loi_, x_LN-INT_* and one can construct similar estimators. The derivation of the respective estimators is based on Eqn. 2.16 from Haviv (20).

### Design of simulations

To assess the performance of our new estimator, we compared it in a simulation study to a selection of other estimators from the literature. Simulations were performed using R 3.5.1 (21) with the prevtoinc package which will be freely available on CRAN (http://cran.r-project.org/).

In a first step, we assessed the quality of the estimators for *x_loi_, x_LN-INT_,x_los_*.

The setup was the following: a distribution for *X_loi_* was chosen and the corresponding distributions for *A_loi_* and *L_loi_* were derived. A sample of *n* values from *A_loi_* and *L_loi_* was drawn and, based on this sample, all considered estimators of *x_loi_* were calculated. We repeated this procedure *m* times and calculated the root-mean square deviation (RMSD) for each estimator.

An analogous procedure was used to benchmark estimators for *x_los_* and *x_LN-INT_*.

We performed repeated simulations to assess the performance of estimators for *I* based on simulated PPS data as follows: The number of patients *n* in the PPS was fixed, as well as a distribution for *X_loi_* and a value *P* = 0.05 was fixed for the prevalence. For each patient, the presence of a HAI was determined by a sample from a Bernoulli distribution with as parameter *P*. In a next step, for patients with HAIs a joint sample of *A_loi_* and *L_loi_* was sampled from the chosen distribution. To assess the performance of the estimators for *I_pp_* we additionally sampled *A_los_* and *L_los_* jointly for all patients. For a simulation distribution of *X_LN-INT_*, assessment of estimators was performed in an analogous way replacing *P* by *P_rhame_* = 0.2. For further parameters of the simulations see supplement S3.

### Estimators for comparison

We used the following estimators to benchmark the performance of our new estimator.

- *pps.median* - estimator based on the median duration up to PPS (1)

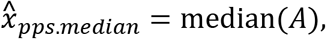

where median(*A*) is the median of samples of the observed *A*,
- *pps.mean* - alternative estimator used in (1) based on the mean instead of the median

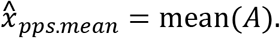
- *L.full* - estimator based on samples from the PPS with information on *L* based on the transformation formula 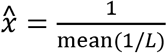

The transformation formula uses the theoretical relationship between *X* and *L* derived in Eqn. 7 in (8). When comparing the performance of estimators, one has to keep in mind that *L.full* uses the information on the whole durations *L* instead of only information on *A*.

The estimators can be used to estimate *x_loi_, x_LN-INT_* or *x_los_* depending on which duration up to PPS *A* we use.

### Mixed estimator

We also experimented with the combination of different estimators by weighting. As will be seen in the results section, for small samples the estimator *gren* has high variance. While it is unbiased (inside the model), it could be advantageous to combine it with a biased estimator with lower variance for small sample sizes. As a specific case, we introduced the following estimator *pps.mixed* based on the estimators *pps.mean* and *gren:*

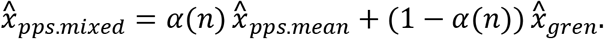

The function *α* is chosen as a sigmoid function: 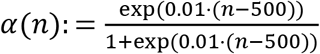. This gives a smooth transition between *pps.mean* and the new estimator *gren* with equal weighting *α* = 0.5 on *n* = 500. Again this type of estimator can be used for the estimation of *x_loi_, x_LN-INT_* or *x_los_*.

### Constructing estimators for *I* and *I_pp_*

We estimated the theoretical prevalence *P* by taking the observed prevalence 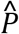 on the day of the PPS as an estimate. We constructed the incidence rate estimator in the general form:

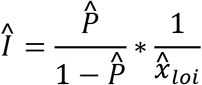

and for the incidence proportion per admission:

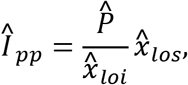

where one uses any of the above estimators for 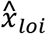 and 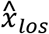. A similar estimator could be built by plugging in the corresponding estimators in the original Rhame-Sudderth formula using *P_rhame_* and *x_LN-INT_*.

### Simulation distributions for *X_loi_, X_LN-INT_* and *X_los_*

We used three different distributions for *X_loi_*: a geometric distribution shifted to start on 1 with mean 8, a Poisson distribution shifted to start on 1 with mean 8. We selected the two theoretical distributions, Poisson and geometric, to assess the flexibility of the estimators. We also used an empirical distribution of *X_loi_* based on data from the NIDEP2 – study (15). In this study, incidence and prevalence of HAIs were measured on a daily basis in eight German hospitals during two eight-week periods (see supplement S2 for a further description of the data).

For simulation of *X_LN-INT_*, we used an empirical distribution of *X_LN-INT_* based on the HAI-Net ICU data from 2015, which monitored date of onset of HAI and date of discharge of patients with an ICU-acquired HAI in 1 365 intensive care units (ICUs) from 11 European Union Member States (see supplement S2 for a further description of the data) (15, 16). No information on the end of the HAI was available, which is why we used *X_LN-INT_* and the original version of the Rhame-Sudderth formula for this simulation example.

We show the resulting distributions of *X_loi_* in Fig. 2 and the distribution for *X_LN-INT_* in Fig. 3.

**Fig. 2.**
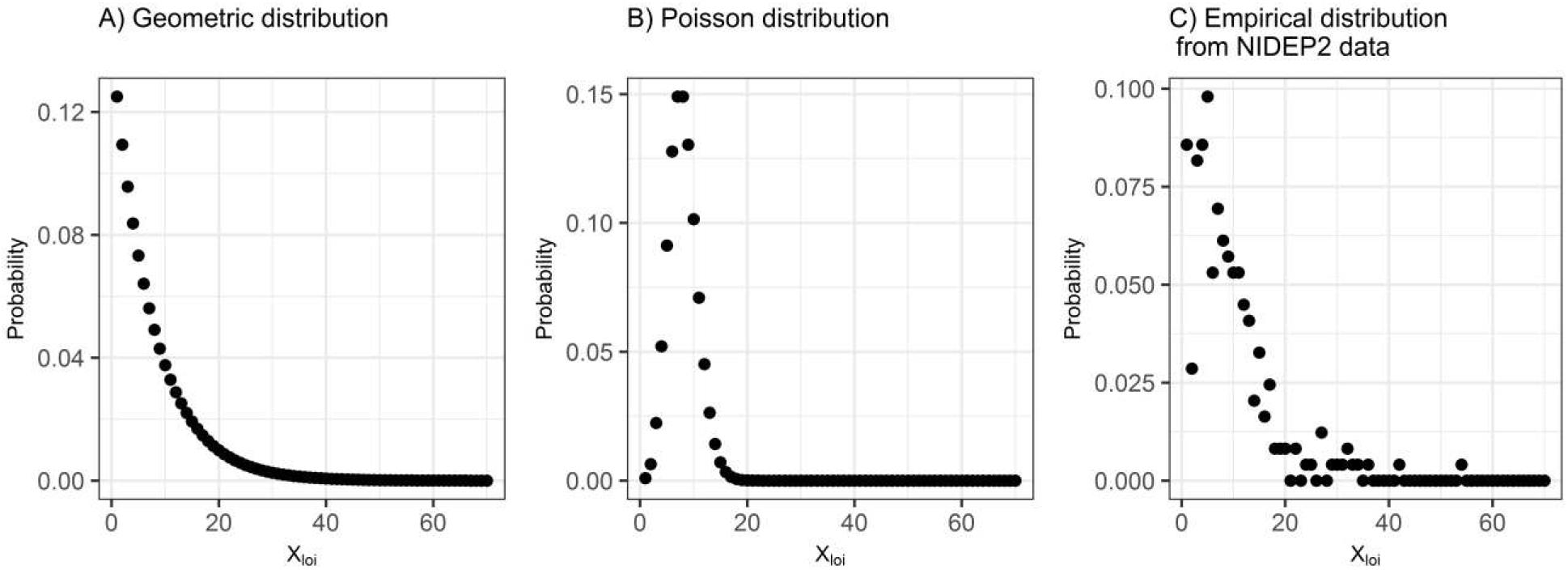
Distributions of X_loi_ for simulations

**Fig. 3.**
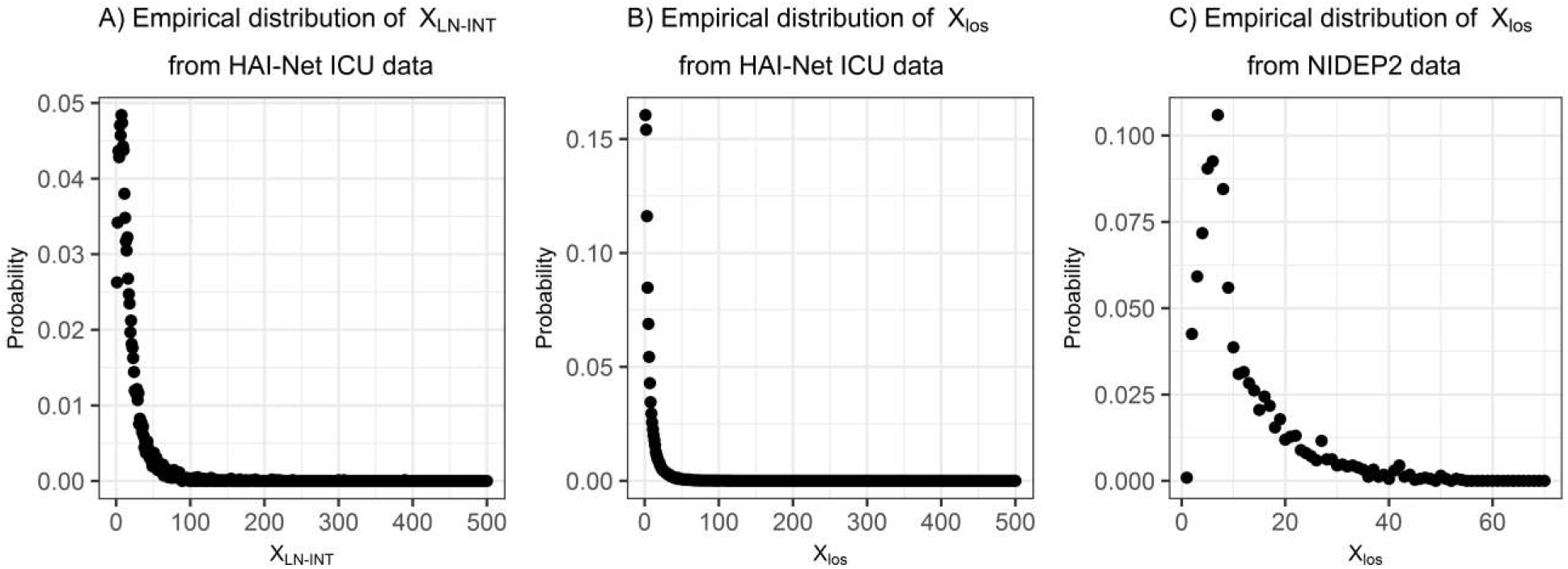
Empirical distribution of X_LN-INT_ from HAI-Net ICU data (A) and empirical distributions for X_los_ for NIDEP2 and HAI-Net ICU data (B and C)

Based on these distributions for *X_loi_* we calculated the distributions of *A_loi_* and *L_loi_* (see Eqn. 2.14 and 2.16 (20) for the exact relation between these distributions).

For each simulation we then sampled *n* lengths of infection jointly from *A_loi_* and *L*_loi_. An analogous procedure was applied for sampling lengths of stay (*A_los_* and *L_los_*) and lengths of stay after infection (*A_LN-INT_* and *L_LN-INT_*). The distributions used for simulating the length of stay are shown in Fig. 3 and were based on data on lengths of stay from the NIDEP2-study (14) and HAI-Net ICU (15, 16).

## RESULTS

To assess the quality of the estimators, we measured the RMSD for increasing numbers of HAIs using different distributions (Fig. 4–7).

**Fig. 4.**
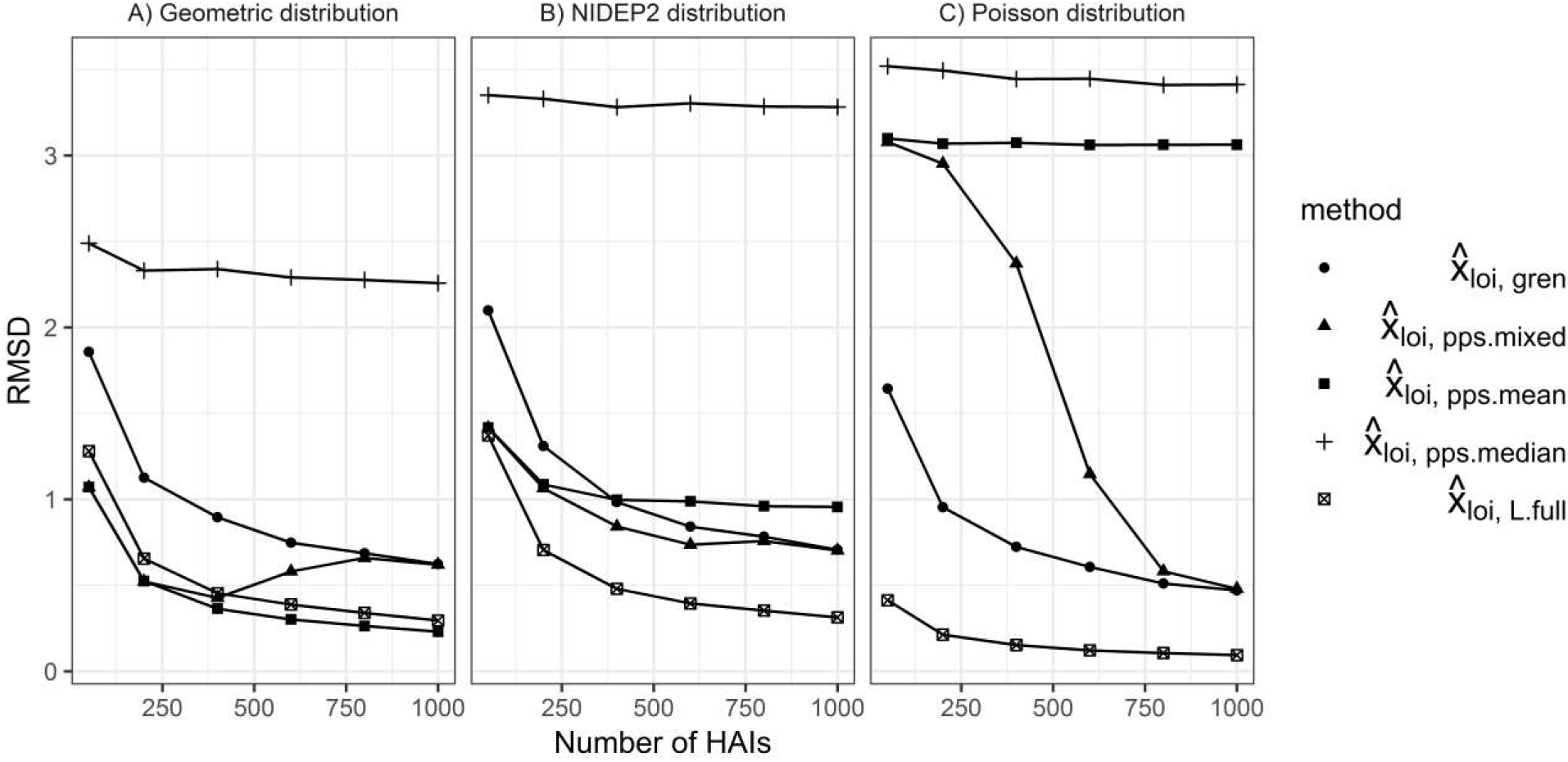
Root-mean squared deviation (RMSD) of estimators of *x_loi_* for 1000 simulations each along increasing size of samples of *A_loi_* (resp. *L_loi_* for L.full)

### Simulations for *x_loi_* and *x_LN-INT_*

In Fig. 4, we present the RMSDs of the estimates of *x_loi_*. We show the results for three examples of *A_loi_* distributions. The simulations ranged from *n* = 50 to *n* = 1000. The estimators differed in the size of the RMSD, as well as in the convergence to zero along increasing sample sizes. In all three distributions, *pps.median* had the highest RMSD and generally did not converge to zero. The estimator pps.mean behaved similarly to *pps.median* in the case of the Poisson distribution. For the NIDEP2 distribution, it did not converge to zero, but stabilized on a lower RMSD compared to the Poisson distribution. In the case of the geometric distribution, *pps.mean* converged to zero with a low RMSD as could be expected for mathematical reasons (22), because *x_loi_* = *a_loi_* for this specific distribution. *L.full* converged towards zero for all distributions and had among the lowest RMSD for all three settings. The RMSD of the new estimator *gren* converged towards zero in all three settings for large enough sample sizes and the magnitude of the RMSD was similar for all three distributions. The RMSD of *pps.mixed* for lower sample sizes was similar to the one of *pps.mean* and for larger sample sizes more like the new *gren* estimator as expected due to its construction.

Similar plots for the bias (inside the model) and standard deviation can be found in the supplement S4 in Fig. S2 and S3. For boxplots of the estimators of *x_loi_x_lnint_* and *x_los_* see Fig. S4–S6 in supplement S5.

Results for the estimation of *x_LN-INT_* for the HAI-Net ICU data are shown in Fig. 5. The simulations again ranged from *n* = 50 to *n* = 1000.

**Fig. 5.**
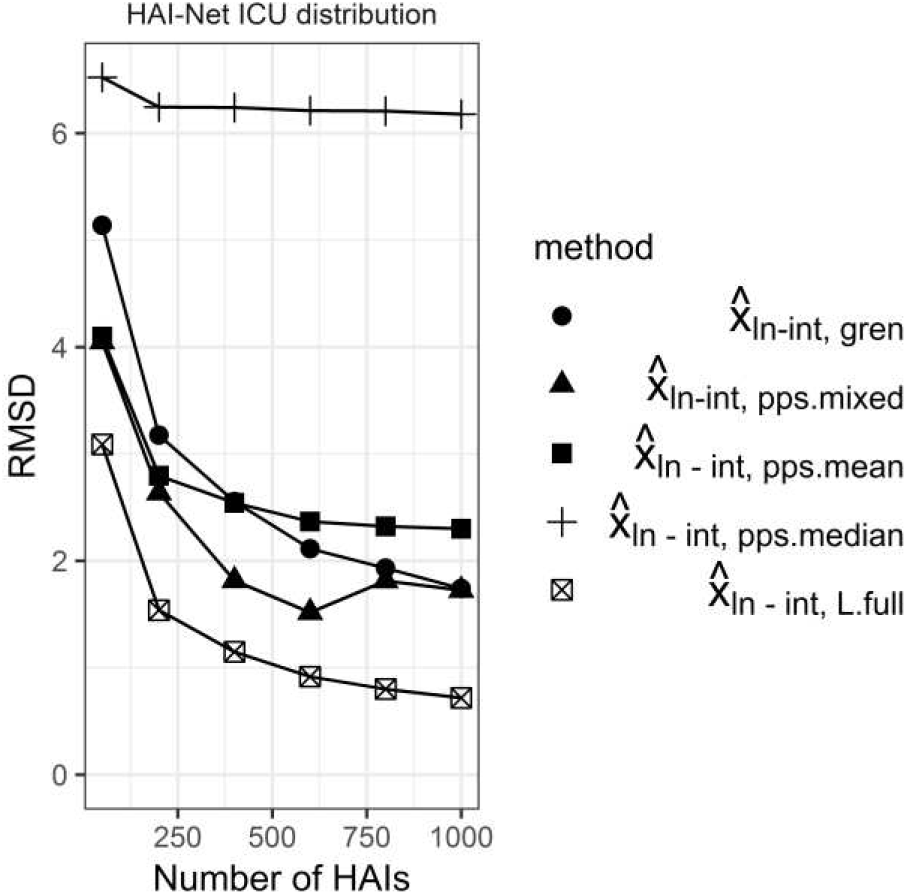
Root-mean squared deviation (RMSD) of estimators of x_LN-INT_ for 1000 simulations each along increasing size of samples of A_LN-INT_ (resp. L_LN-INT_ for L.full)

As previously, *pps.median* did not converge to zero and had the highest RMSD among the estimators. The estimator *pps.mean* stabilized at a significantly lower RMSD than *pps.median*, but did not converge towards zero either. *L.full* was again the best performing estimator in terms of RMSD. *gren* and *pps.mixed* behaved similarly to the estimation of *x_loi_*. *gren* exhibited a lower RMSD than *pps.mean* as the sample size increased, and the RMSD behaviour of *pps.mixed* with increasing sample size was between that of *pps.mean* and *gren*.

### Simulations for *I*

The results for the RMSDs of the estimators for *I* are shown in Fig. 6. In this figure the RMSD was divided by the theoretical incidence rate *I* to estimate the relative size of the error. The sample sizes ranged from *n* = 500 to *n* = 20000 patients in the simulated PPS. As expected, the RMSDs behave very similarly to the case of estimation of *x_loi_* and *x_LN-INT_* and the additional uncertainty in the estimation of *P* did not change the general patterns for the RMSDs shown in Fig. 4 and Fig. 5 when compared to Fig. 6.

**Fig. 6.**
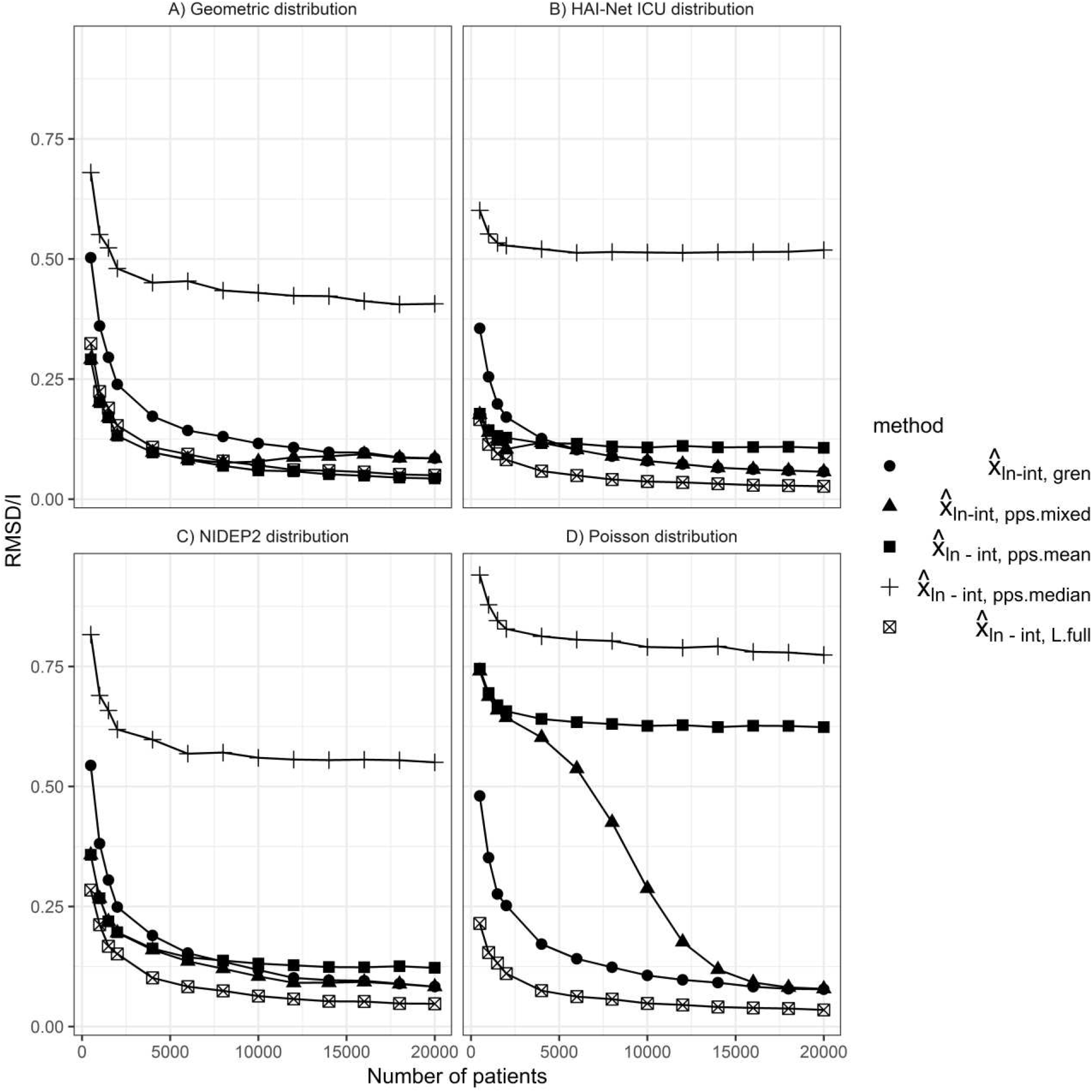
Root-mean squared deviation (RMSD) of estimators of I divided by theoretical incidence rate I along increasing size of samples from a simulated PPS based on 1000 simulations

### Simulations for *x_los_*

Results for the length of stay in days are shown in Fig. 7. Again we presented the RMSD of the estimators of *x_los_*. We used the empirical distributions of the length of stay from the NIDEP2 and HAI-Net ICU datasets.

**Fig. 7.**
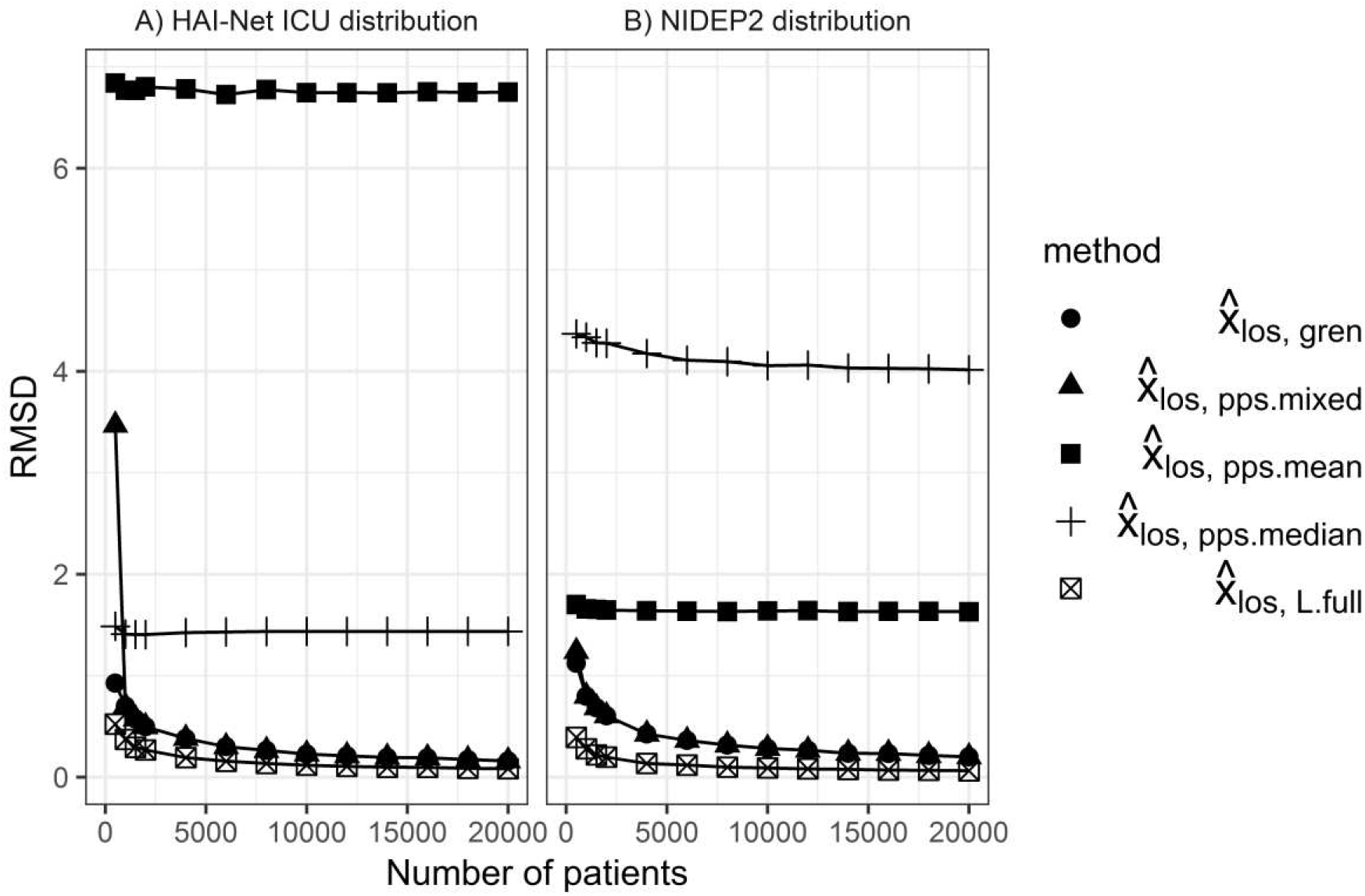
Root-mean squared deviation (RMSD) of estimators of x_los_ using 1000 simulations each along increasing size of samples of A_los_

For the NIDEP2 distribution of lengths of stay, *pps.median* again had the highest RMSD and did not converge toward zero. *pps.mean* did not converge toward zero either but stabilized at a lower RMSD than *pps.median. L.full* again had the lowest RMSD and *pps.mixed* and *gren* also had comparably low RMSD for larger samples (*n* ≥ 5000).

For the HAI-Net ICU distribution of length of stay, the previous picture with respect to *pps.mean* and *pps.median* was reversed. *pps.mean* had the highest RMSD and did not converge towards zero, and *pps.median* had a lower RMSD but also did not converge towards zero. For *gren* and *L.full* the simulation results were similar to those obtained with the NIDEP2 distribution of length of stay. The estimator *pps.mixed* had a high RMSD compared to the other estimators for small sample sizes where the *pps.mean* component was dominant. For larger sample sizes, it behaved similarly to *L.full* and *gren*.

### Simulations for *I_pp_*

For the incidence proportion of HAIs counted per admission, the RMSDs of the estimators are shown in Fig. 8 In this figure the RMSD was divided by the theoretical incidence proportion *I_pp_* to estimate the relative size of the error. Almost all the estimators behaved very similarly in terms of RMSD. *pps.mean* and *pps.median* were the only estimators with a significantly higher RMSD than the other estimators for the HAI-Net ICU distribution. In the case of the NIDEP2 distribution and *pps.median*, the errors in the estimation of *x_loi_* and *x_los_* seemed to cancel out almost exactly, reducing the RMSD for *I_pp_* to levels comparable to the other estimators.

**Fig. 8.**
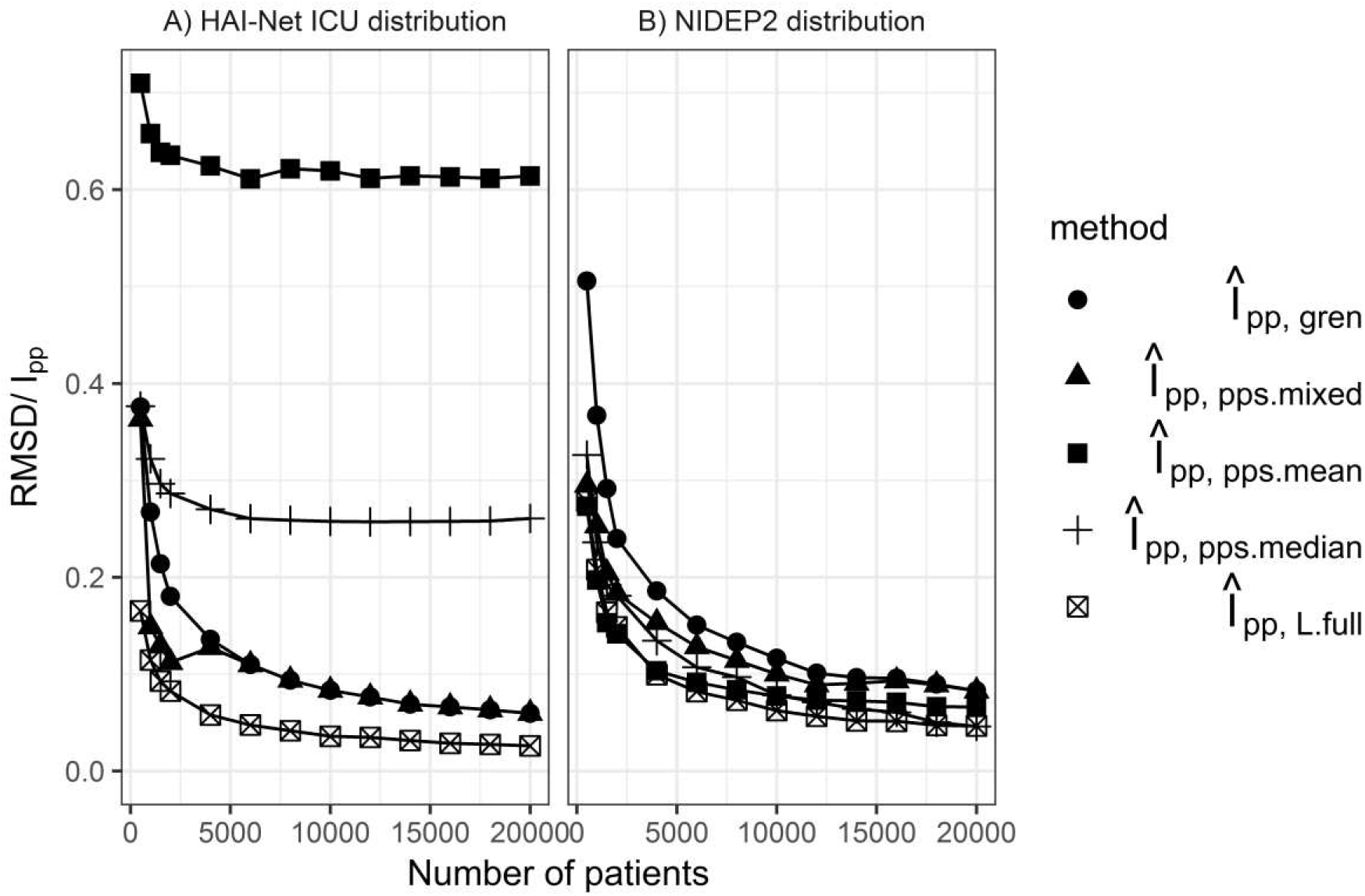
Root-mean squared deviation (RMSD) of estimators of I_pp_ divided by theoretical incidence proportion I_pp_ per admission along growing number of samples from a simulated PPS based on 1000 simulations

## DISCUSSION

We presented a method to estimate incidence from prevalence data available in a typical PPS setup. We used nonparametric estimators for length of stay and length of infection which exploit the monotonicity of *A_los_* and *A_loi_*. By means of a simulation study, we compared these estimators to other estimators that have been applied in previous studies.

The new *gren* estimator behaved consistently for different distributions, i.e. was more accurate with larger samples and in most cases was comparable to or better than the other estimators based on *A*. This was in contrast to *pps.median*, which generally did not converge to the true value. The estimator *pps.mean* did not perform overall as well as the *gren* estimator, but its variance for small sample sizes was lower. As expected, *L.full* performed better than or as well as all other estimators across all settings, but at the price of requiring knowledge of the full durations *L* which are typically not available in a PPS. We finally proposed the mixed estimator *pps.mixed* as a good compromise between the low variability of the *pps.mean* for smaller samples and the consistent behaviour of the new estimator *gren* for larger samples. Altogether, the incidence estimate based on a PPS can be improved by more than 40% of the theoretical value (in terms of RMSD), compared to other estimators from the literature for a large enough PPS (see Fig. 8).

The new method presented in this article is a modification and update of the Rhame-Sudderth formula and is applicable in the setup of modern PPSs. The Rhame-Sudderth formula was published in the 1980s and, to our knowledge there have only been few methodological contribution addressing the questions of validity of the formula on a theoretical level since its publication. Mandel and Fluss (23) have proposed and studied incidence estimators, which generalize the Rhame-Sudderth, estimator but they depend on the use of the original Rhame-Sudderth prevalence definition and information about the total length of stay *L_los_* for all patients in the survey. There have been attempts to evaluate the Rhame-Sudderth formula (14, 24), but these shared the limitation of using *P* instead of *P_rhame_*, as intended in the original Rhame-Sudderth formula. Few studies distinguish between *P* and *P_rhame_* and use the originally intended combination of prevalence and length of duration definitions. This often leads to the use of *x_LN-INT_* as a proxy for length of infection *x_loi_* (14, 24). It was suggested that *x_LN-INT_* was not a good proxy for average length of infection (13) and instead some ad-hoc measure or external information could be used to estimate average length of infection (9, 11–13). Most of the articles remained critical of their own results. The ECDC-coordinated PPS of healthcare-associated infections and antimicrobial use in European acute care hospitals included data from over 200 000 patients across Europe and used the estimators *pps.mean* and *pps.median* (or more precisely a combination of these two) to estimate *x_LN-INT_* stratified by participating country (1). Information on *x_los_* was often obtained from external data sources. In the analysis of the latest ECDC-coordinated PPSs in acute care hospitals and long-term care facilities (2016-2017) (14) our proposed method has already been used for sensitivity analysis to compare with the estimator described above. European-level estimates were similar for the different estimators with few exceptions at individual country level.

The United States PPS coordinated by CDC (3) used stratification along factors thought to be predictive of the prevalence of HAIs. The estimators in (2) were based on medians of the durations-up-to-PPS similar to *pps.median* or external information and in (4) the original Rhame-Sudderth formula was used with the definition of prevalence *P* instead of *P_rhame_* and with a length-biased version of *x_LN-INT_* (i.e. *l_LN-INT_*), and a length-biased version of *x_los_*(i.e. *l_los_*).

A main strength of our method is that we do not make any assumptions on the distributions of *X_loi_* and *X_los_*. Usually in a PPS we do not know the distribution of *X_loi_* and *X_los_*. This means that one criterion for selecting an estimator of *I* or *I_pp_* is that it should behave well irrespective of the form of the unknown distribution. This is a criterion which, among the estimators using only duration-up-to-PPS information, was only fulfilled be the proposed estimator *gren* and the mixed estimator *pps.mixed* for larger samples (*n* ≥ 500). This is supported not only by simulations, but also by theoretical considerations. Using only simulation studies to assess an estimator can be a source of error if the distributions on which the estimators are assessed differ significantly from the underlying distributions encountered in PPSs.

Our method has limitations. One is the requirement of a sufficiently large sample size to get an acceptable estimate. We took the sample size of 500 HAIs as a rule-of-thumb lower limit. It may be applied to smaller samples, but with a risk of lower precision. For a single medium-size hospital, repeated PPSs with aggregation of the results would need to be performed to reliably estimate the incidence of HAIs. Another limitation of our setup is that we counted multiple simultaneous or partially overlapping HAIs as one HAI. However, these in reality only comprise a small fraction of HAIs (1, 15) and therefore can be neglected. In addition, many of the limitations mentioned in the original article by Rhame and Sudderth also apply in this updated version, in particular the lack of explicit representation of outbreaks and the assumption that the risk that a patient acquires a HAI is independent of other patients’ status. The new estimators *gren* and *pps.mixed* applied to the length of stay are sensitive to week day patterns in admissions and discharges (data not shown).

Typically, for larger PPSs, data collection takes place on different weekdays for different hospitals or even different wards in the same hospital (1), thus mitigating the influence of these patterns on the estimates. Another issue is that patients on their first day of admission are sometimes underrepresented due to the PPS protocol, when e. g. only patients admitted before a fixed time are included in the PPS. The new estimators are based on the monotonicity assumption for the distribution of *A_los_*, which is violated in this situation. One solution can be to let *A* denote full days of hospital stay and ignore the patients admitted on the date of the survey for the estimates of average length of stay, but include them in the estimate of the prevalence. Similar problems appear to a lesser extent for the first day of HAI. Other factors that need to be taken into account include the consistency of the application of case definitions for HAIs, and the representativeness of the hospital sample.

In conclusion, the proposed *gren* estimator and the combined estimator *pps.mixed* provide better estimates of the length of infection across a range of simulation settings when compared to previously used estimators and, in contrast to these, are grounded in theory. The simulations also serve as a guide of the sample size to include in a PPS required to estimate incidence. The method is shared and easily applicable with the help of the R package `prevtoinc`.

## Abbreviations

CDC–: Centers for Disease Control and Prevention
ECDC–: European for Disease Control and Prevention
HAI–: healthcare-associated infection
HAI-Net ICU–: European surveillance of HAIs in intensive care units
NIDEP2–: second study of nosocomial infections in Germany (Nosokomiale Infektionen in Deutschland, Erfassung und Prävention)
PPS–: point-prevalence survey RMSD - root mean squared deviation

## Supplement: From prevalence to incidence - a new approach in the hospital setting

### S1: Mathematical model and technical details

#### Derivation of the conversion formula

The theoretical model used is that of a discrete renewal process (see Chapter 2, [1] for an introduction to renewal theory). Similar to Rhame and Sudderth [2] we see a single bed as the basic unit to be simulated. Beds are assumed to be occupied sequentially by patients, which can develop HAIs on each day of stay. The evolution of patients/infections per bed are assumed to be statistically independent. We assume that time *t* is progressing in patient-days.

We use the framework of an alternating renewal process between HAI / non-HAI. Theorem 4.8 from [3] gives us the relation

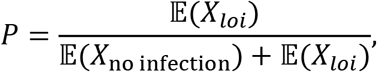

where *X*_no infection_ is the length of occupation of a bed without any infection (the time in between two infections) and E will denote the expected value in the following. In our model we assume that the probability of acquiring a HAI on a given day is *I* and therefore one can see that in our model *X*_no infection_ follows a geometric distribution with parameter *I*. This means 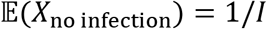. It follows:

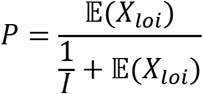

After rearrangement this gives:

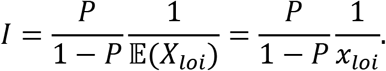

To get the version per patient we first introduce the notation *X*_los w/o HAI_ for the length of stay of a patient only counting days without a HAI. We can then write:

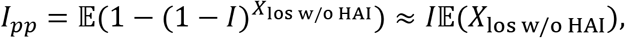

where the approximation is valid if 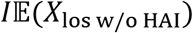 is small.

Finally, we note under the assumption that *X_los_* is independent of whether or not an infection occured:

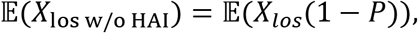

as (1 − *P*) represents the average proportion of time of stay without a HAI present.

A proof of this is based on the following equalities, where for the distribution of limits Slutsky’s theorem is used:

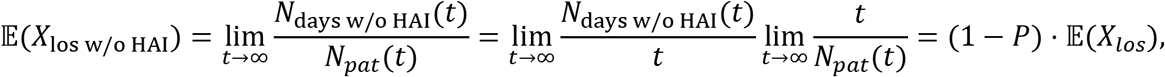

where *N_pat_*(*t*) is the number of patients in the bed up to time *t* and *N*_days w/o HAI_(*t*) denotes the number of patients days without HAI in the bed up to time *t*.

Putting together the above results gives the final conversion formula:

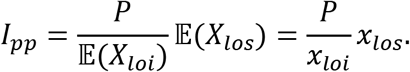

All the theoretical arguments can be repeated by replacing *X_loi_* by *X_LN-INT_* and this would lead to the original Rhame-Sudderth formula. We note that *P* is then defined as the proportion of time a bed is occupied by a person who had or has a HAI, which is the definition of *P_rhame_* from the main text.

#### Asymptotics of *gren* estimators

In our setting the *gren* estimators of the average durations will exhibit asymptotic normality. The proofs are based on the delta method and Prop. 3.4 and Prop. 3.6 from [4]. One gets

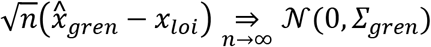

with 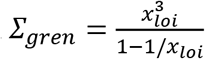 where ⇒ denotes convergence in distribution, *n* the sample size and 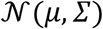 denotes a normal distribution with mean *μ* and covariance *Σ*.

As pointed out in [4] the asymptotics of the Grenander estimator for a discrete distribution are the same as for one based on the empirical estimator, i. e. taking just the empirically observed proportions as an estimate for the distribution of *A*. To visualize the relation between asymptotic results and simulations, we take *x_loi_* as an example. We call the estimator based on the empirical proportions 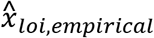. In Fig. S1 we compare the simulated standard deviations for 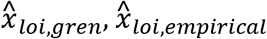 and the asymptotical approximation of the standard deviation 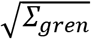. As predicted by theory, the standard deviations of the estimators approach the asymptotic value and also 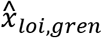 systematically has the lowest standard deviation.

**Fig. S1.**
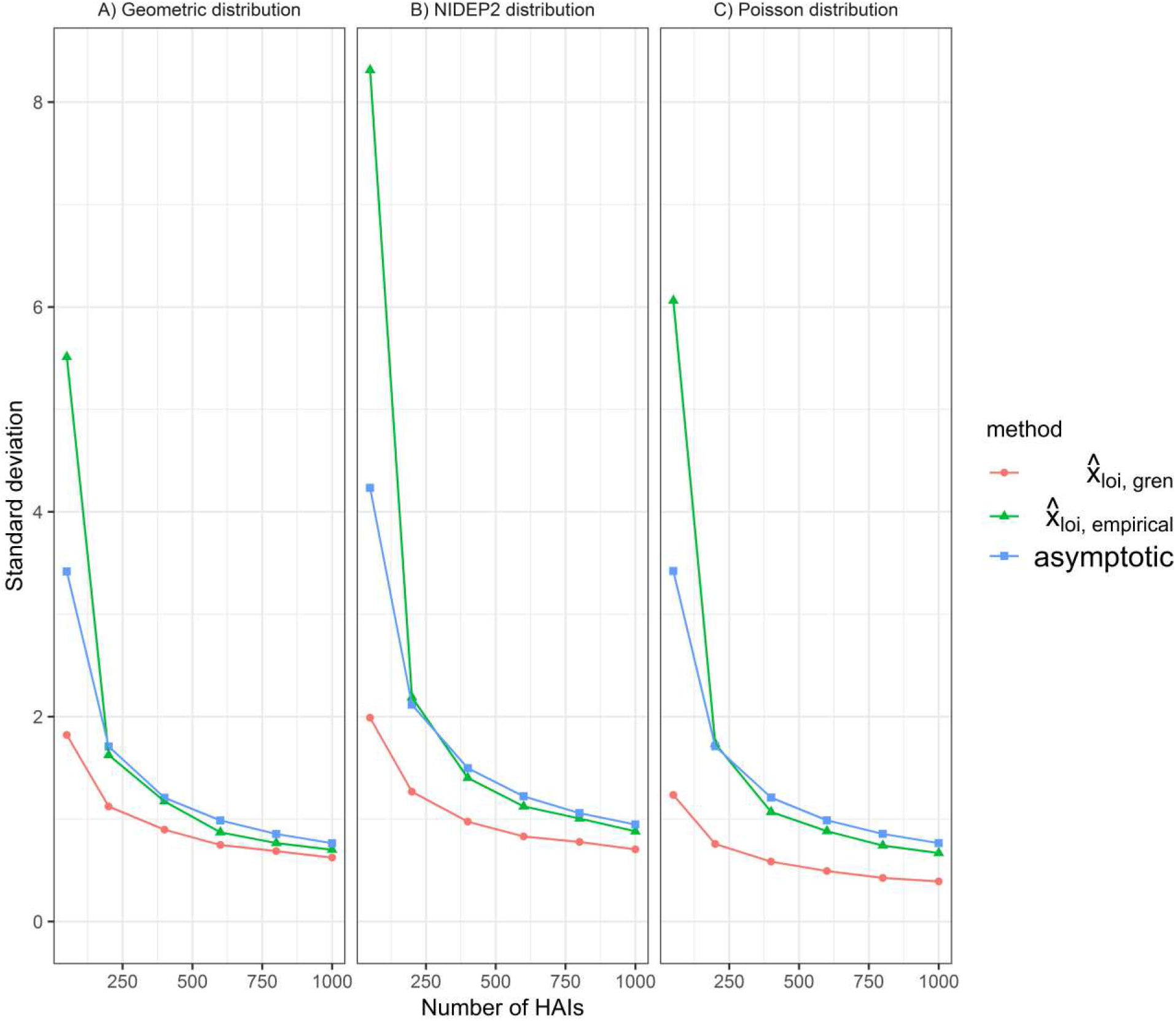
Standard deviation of estimators of for 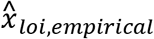 and 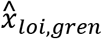 based on 1000 simulations each along growing number of samples of *A_loi_* compared to approximation of standard deviation based on asymptotics (in green)

### S2: Description and characteristics of data sources

#### NIDEP2

As a data source for our simulations we used the NIDEP2 study [5]. We used data from eight German hospitals, where incidence and prevalence of HAIs were measured on a daily basis during two eight-week periods. The second monitoring period began after a randomly assigned intervention to four of the hospitals in which a range of additional infection prevention and control measures was introduced. A total of 7568 patients participated in the NIDEP2 study, among whom 487 had at least one HAI. We counted multiple simultaneous or partially overlapping periods of HAI as one HAI. For the generation of the empirical NIDEP2-distribution of *X_loi_*, we used the incidence data up to day 30 (*n_hai_* = 245) for each period (as this corresponded to the point where the influence of the cutoff of the measurement after each incidence period was not discernible anymore). Infections already present on the first day of each period were excluded because there was no information on their duration before the start of the surveillance period.

#### HAI-Net ICU

As a second data source we used data from the European surveillance of HAIs in intensive care units (HAI-Net ICU) [6,7]. In 2015, a total of 141 955 patients from 1 365 intensive care units (ICUs) from 11 European Union Member States were included. Among these patients, 11 788 developed at least one HAI during their stay [7]. In the HAI-Net ICU dataset, the onset of the HAI and the date of discharge were available, but not any information on the end of the HAI, so we calculated *X_LN-INT_* and used the original version of the Rhame-Sudderth formula to calculate estimates of *x_LN-INT_*. This meant that we also used *P_rhame_* as the measure of prevalence as we could not estimate *P* without the information at the end of the HAI. Based on the ICU data, *P_rname_* was around 22% for most of the year with a dip around the beginning/end of the year, which could be due to reporting practices. A further modification was to discount the first two days of stay in the ICU. According to the HAI-Net ICU protocol, infections are only considered ICU-acquired if they develop at least 48 hours after admission to the ICU [6]. All the estimators were modified accordingly.

### S3: Specification of simulation parameters

We measured the performance of the methods by repeating the simulation 1000 times for different values of the sample size n. With this we estimated the bias (inside the model), standard deviation and RMSD of the estimators for *x_loi_* and for the case of the HAI-Net ICU distribution *x_LN-INT_*. The theoretical values for *x_loi_* were the following: *x_ioi,geom_* = 8 for the geometric distribution, *x_loi,nidep2_* = 9.28 and *x_loi,pois_* = 8. In a next step, we assessed the performance of the methods for estimating I. We used a setup with a theoretical *P* = 0.05. This meant that each simulated patient had a probability of 0.05 having an active HAI on the day of the PPS. Taking this theoretical prevalence as given, we generated simulated PPS data with associated *L_loi_* and *A_loi_* for each patient with a HAI. For the HAI-Net ICU data, the same simulations were performed with *X_LN-INT_* and *P_rname_* = 0.20 in place of *X_loi_* and *P*. The average number of days from first HAI to discharge in the distribution was *x_LN-INT,icu_* = 18.38. The theoretical incidence rate in the simulated models was *I* = 0.0066 (6.6 HAIs per 1000 patient-days at risk) for the Poisson and geometric distribution, *I* = 0.0066 (6.6 HAIs per 1000 patient-days at risk), *I* = 0.0057 (5.7 HAIs per 1000 patient-days at risk) for the NIDEP2 distribution and *I* = 0.0136 (13.6 HAIs per 1000 patient-days at risk) for the HAI-Net ICU distribution. These incidence figures should not be confused with the incidence that could be calculated from the original data sets. The results that we used were based on a fictitious fixed prevalence used for the simulations. We repeated these simulations along increasing sample sizes (numbers of patients) *n* with 500 simulations for each *n* to estimate the RMSD. To assess the quality of estimation of the length of stays, we also performed a simulation based on the empirical distribution of lengths of stay in the NIDEP2 and HAI-Net ICU datasets. The procedure was analogous to the case of the length of infection. The mean values of the length of stay for the two distributions were: *x_los,icu_* = 8-44 (discounting the first two days) and *x_los,nidep2_* = 11-01. Finally, we assessed the estimators for *I_pp_* by combining the distributions of length of stay and length of infection (or length of stay after infection) based on the NIDEP2 and HAI-Net ICU data under the assumption that the product of the marginal distributions of these quantities is a good approximation of the joint distribution. We used the same parameters and simulation sizes as for the estimation of I; the theoretical *I_pp_* = 0.062 for the simulations from NIDEP2 data and *I_pp_* = 0.027 from HAI-Net ICU data.

### S4: Bias and standard deviation of the estimators for *x_loi_*

**Fig. S2.**
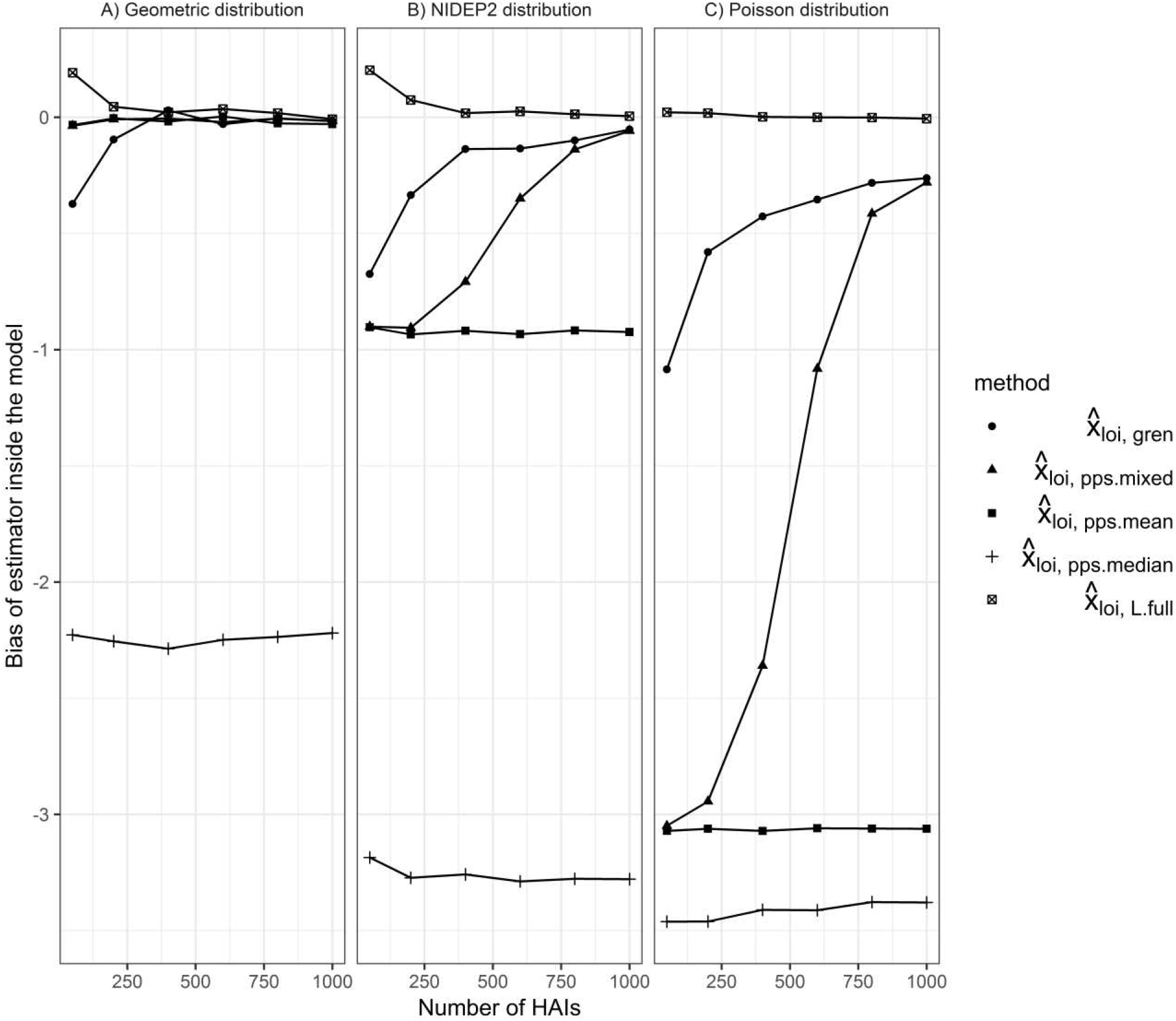
Bias of estimators (inside the model) of x_loi_ for 1000 simulations each along growing number of samples of A_loi_

**Fig. S3.**
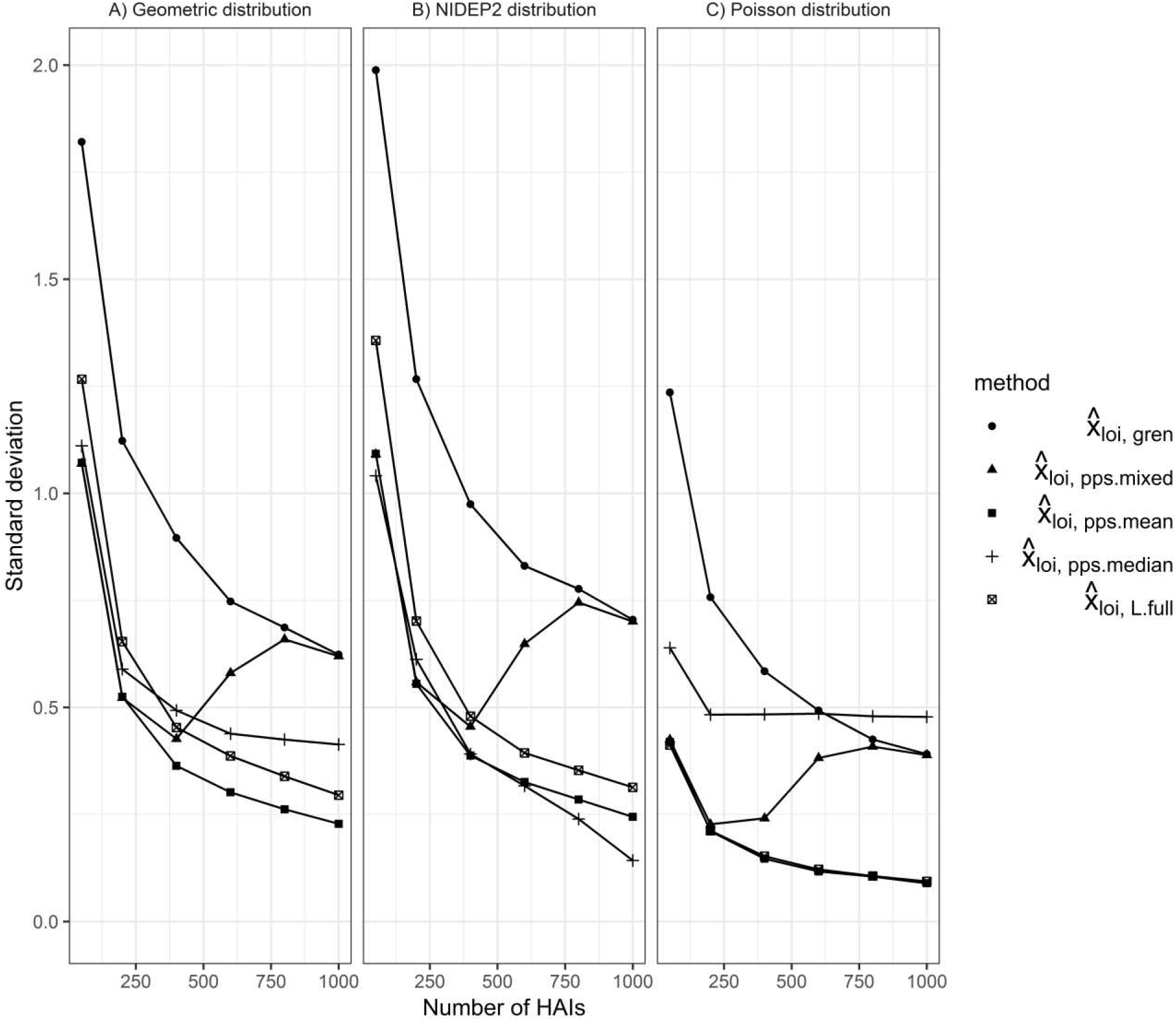
Standard deviation of estimators of x_loi_ for 1000 simulations each along growing number of samples of A_loi_

### S5: Boxplots for simulations of estimates for *x_loi_, x_LN-INT_* and *x_los_*

In this section, we present boxplots of the estimates for *x_loi_, x_LN-INT_* and *x_los_*. The boxplots are specified in the following way: The lower and upper hinges correspond to the first and third quartiles. The upper whisker extends from the hinge to the largest value no further than 1.5 · IQR away from the hinge (IQR being the inter-quartile range). The lower whisker extends from the hinge to the smallest value at most 1.5 · IQR away from the hinge. Data points beyond the end of the whiskers are plotted individually.

**Fig. S4.**
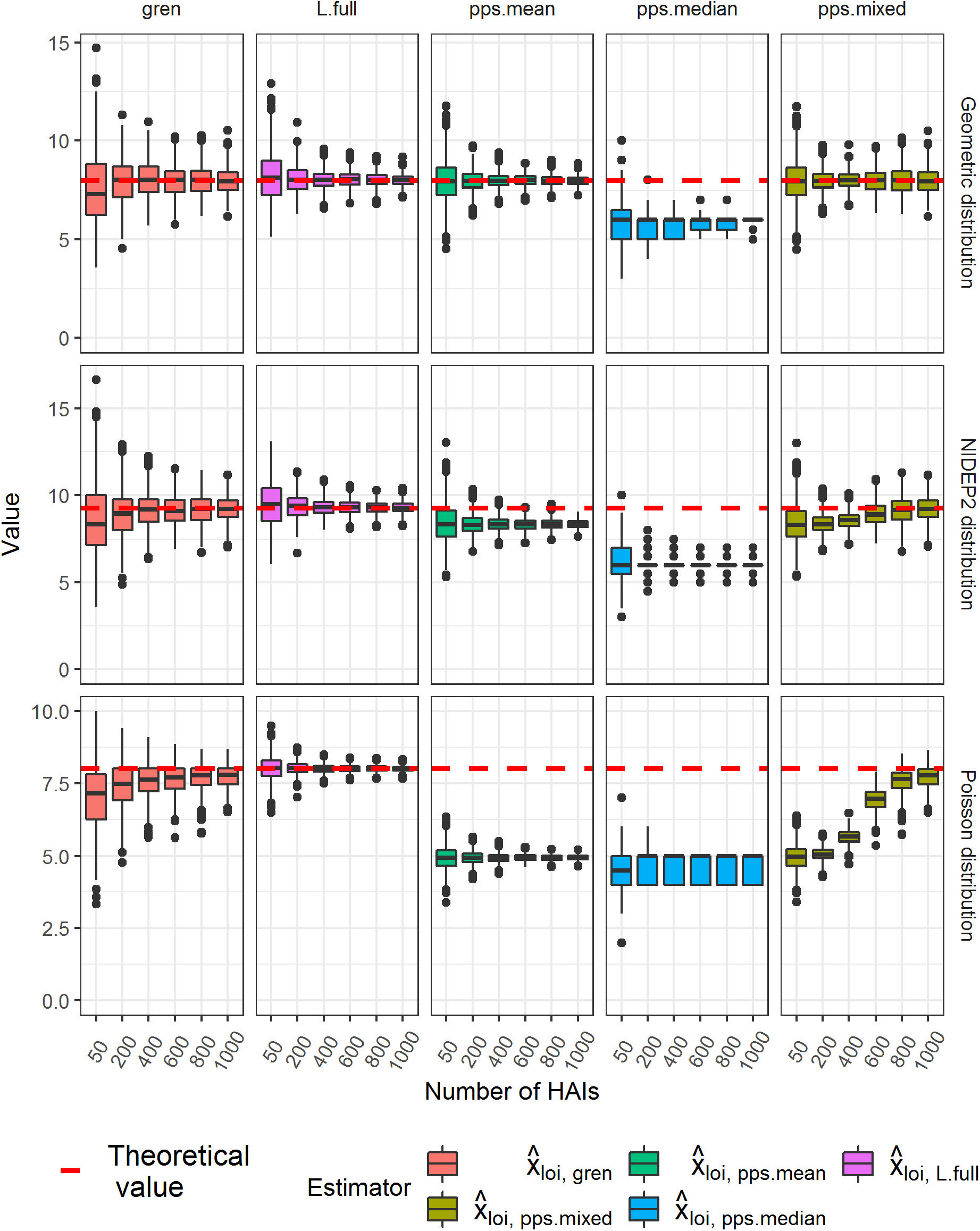
Boxplots of estimates of x_loi_ for 1000 simulations each along growing number of samples of A_loi_

**Fig. S5.**
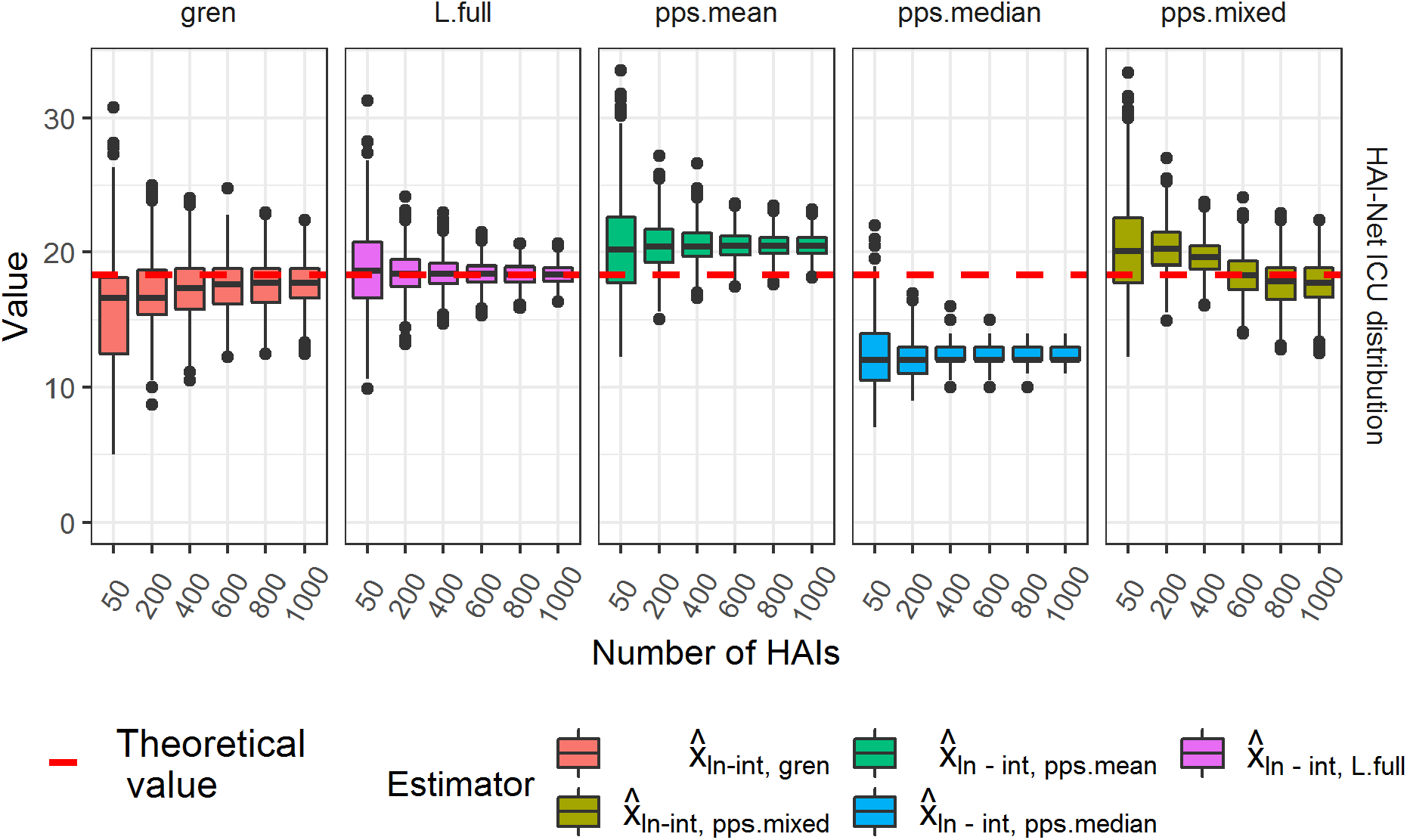
Boxplots of estimates of for x_LN-INT_ for 1000 simulations each along growing number of samples of A_LN-INT_

**Fig. S6.**
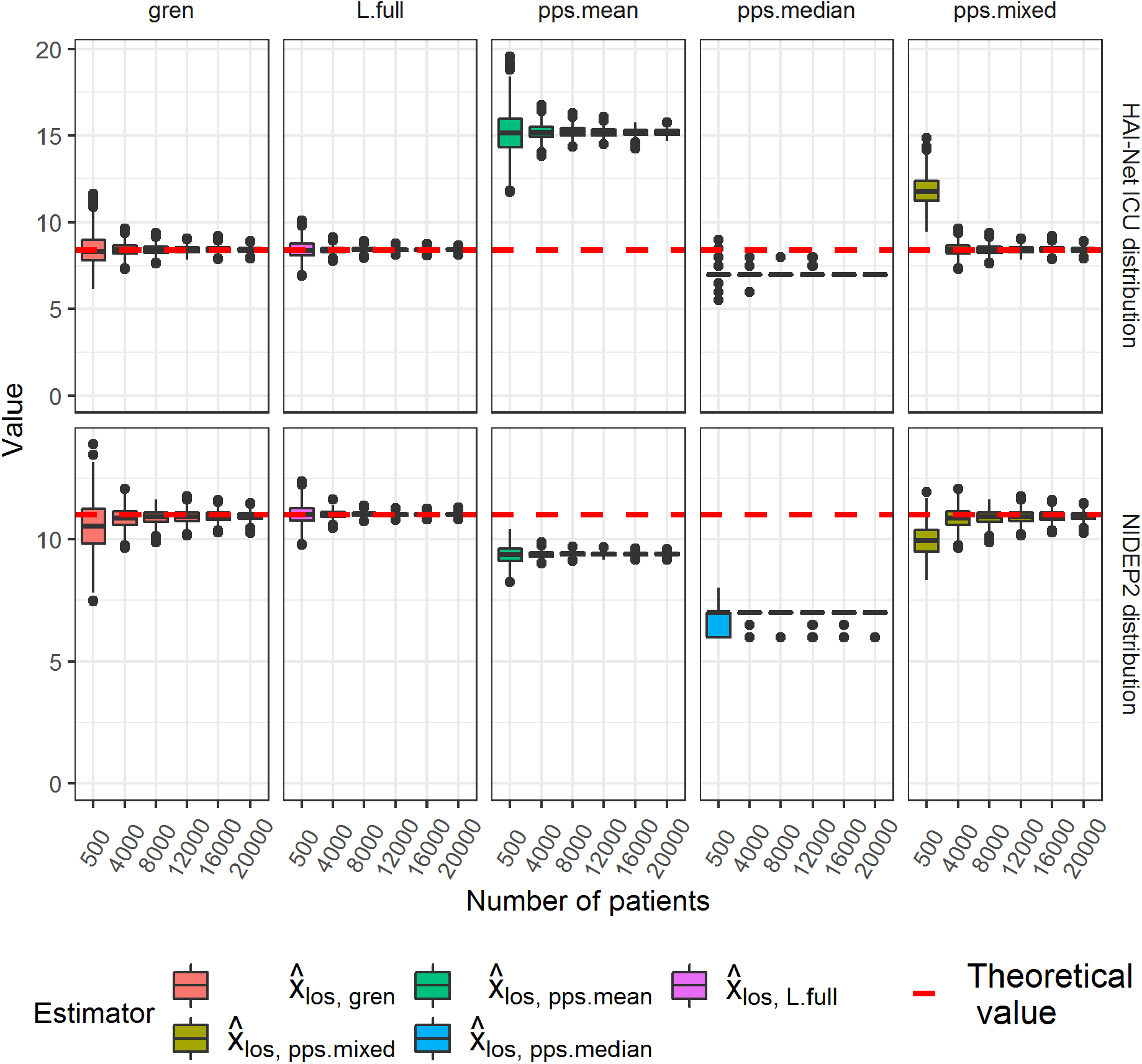
Boxplot of estimates of x_los_ for 1000 simulations each along growing number of samples of A_los_

## References

1. European Centre for Disease Prevention and Control. Point prevalence survey of healthcare-associated infections and antimicrobial use in European acute care hospitals. ECDC, Stockholm. 2013. http://ecdc.europa.eu/en/publications/Publications/healthcare-associated-infections-antimicrobial-use-PPS.pdf.

2. Suetens C, Latour K, Karki T, et al. Prevalence of healthcare-associated infections, estimated incidence and composite antimicrobial resistance index in acute care hospitals and long-term care facilities: results from two European point prevalence surveys, 2016 to 2017. Euro Surveill. 2018;23(46). doi:10.2807/1560-7917.ES.2018.23.46.1800516

3. Magill SS, Edwards JR, Bamberg W, et al. Multistate point-prevalence survey of health care-associated infections. N Engl J Med. 2014;370(13):1198–208. doi:10.1056/NEJMoa1306801

4. Magill SS, O’Leary E, Janelle SJ, et al. Changes in Prevalence of Health Care-Associated Infections in U.S. Hospitals. N Engl J Med. 2018;379(18):1732–44. doi:10.1056/NEJMoa1801550

5. Cassini A, Plachouras D, Eckmanns T, et al. Burden of Six Healthcare-Associated Infections on European Population Health: Estimating Incidence-Based Disability-Adjusted Life Years through a Population Prevalence-Based Modelling Study. PLoS Med. 2016;13(10):e1002150. doi:10.1371/journal.pmed.1002150

6. Rothman KJ, Greenland S, Lash TL. Modern epidemiology. 3rd ed. ed. Philadelphia, Pa.: Lippincott Williams & Wilkins; 2008.

7. Rhame FS, Sudderth WD. Incidence and prevalence as used in the analysis of the occurrence of nosocomial infections. Am J Epidemiol. 1981;113(1):1–11.

8. Freeman J, Hutchison GB. Prevalence, incidence and duration. Am J Epidemiol. 1980;112(5):707–23.

9. Berthelot P, Garnier M, Fascia P, et al. Conversion of prevalence survey data on nosocomial infections to incidence estimates: a simplified tool for surveillance? Infect Control Hosp Epidemiol. 2007;28(5):633–6. doi:10.1086/513536

10. Gastmeier P, Brauer H, Sohr D, et al. Converting incidence and prevalence data of nosocomial infections: results from eight hospitals. Infect Control Hosp Epidemiol. 2001;22(1):31–4. doi:10.1086/501821

11. Graves N, Nicholls TM, Wong CG, Morris AJ. The prevalence and estimates of the cumulative incidence of hospital-acquired infections among patients admitted to Auckland District Health Board Hospitals in New Zealand. Infect Control Hosp Epidemiol. 2003;24(1):56–61. doi:10.1086/502116

12. Kanerva M, Ollgren J, Virtanen MJ, Lyytikainen O, Prevalence Survey Study G. Estimating the annual burden of health care-associated infections in Finnish adult acute care hospitals. Am J Infect Control. 2009;37(3):227–30. doi:10.1016/j.ajic.2008.07.004

13. King C, Aylin P, Holmes A. Converting incidence and prevalence data: an update to the rule. Infect Control Hosp Epidemiol. 2014;35(11):1432–3. doi:10.1086/678435

14. Meijs AP, Ferreira JA, Sc DEG, Vos MC, Koek MB. Incidence of surgical site infections cannot be derived reliably from point prevalence survey data in Dutch hospitals. Epidemiol Infect. 2017;145(5):970–80. doi:10.1017/S0950268816003162

15. Gastmeier P, Brauer H, Forster D, Dietz E, Daschner F, Ruden H. A quality management project in 8 selected hospitals to reduce nosocomial infections: a prospective, controlled study. Infect Control Hosp Epidemiol. 2002;23(2):91–7. doi:10.1086/502013

16. European Centre for Disease Prevention and Control. European surveillance of healthcare-associated infections in intensive care units - HAI-Net ICU protocol, version 1.02. Stockholm. 2015. http://ecdc.europa.eu/en/publications/Publications/healthcare-associated-infections-HAI-ICU-protocol.pdf

17. European Centre for Disease Prevention and Control. Healthcare-associated infections acquired in intensive care units. In: ECDC. Annual epidemiological report for 2015. ECDC, Stockholm. 2017. https://ecdc.europa.eu/sites/portal/files/documents/AER_for_2015-healthcare-associated-infections_0.pdf.

18. Arratia R, Goldstein L, Kochman F. Size bias for one and all. In: ArXiv E-Prints. 2013. https://arxiv.org/abs/1308.2729.

19. Jankowski HK, Wellner JA. Estimation of a discrete monotone distribution. Electron J Stat. 2009;3:1567–605. doi:10.1214/09-EJS526

20. Haviv M. Queues - a course in queuing theory. New York: Springer; 2013.

21. R Development Core Team. R: A language and environment for statistical computing. R Foundation for Statistical Computing, Vienna, Austria; 2018.

22. Freeman J, McGowan JE, Jr. Day-specific incidence of nosocomial infection estimated from a prevalence survey. Am J Epidemiol. 1981;114(6):888–901.

23. Mandel M, Fluss R. Nonparametric estimation of the probability of illness in the illness-death model under cross-sectional sampling. Biometrika. 2009;96(4):861–72. doi:10.1093/biomet/asp046

24. Rossello-Urgell J, Rodriguez-Pla A. Behavior of cross-sectional surveys in the hospital setting: a simulation model. Infect Control Hosp Epidemiol. 2005;26(4):362–8. doi:10.1086/502553

## References

1. Haviv M. Queues - a course in queueing theory. Springer; 2013.

2. Rhame FS, Sudderth WD. Incidence and prevalence as used in the analysis of the occurence of nosocomial infections. American Journal of Epidemiology. 1981;113:1–11.

3. Beichelt F, Fatti P. Stochastic processes and their applications. CRC Press; 2002.

4. Jankowski HK, Wellner JA. Estimation of a discrete monotone distribution. Electron J Statist. 2009;3:1567–605.

5. Gastmeier P, Bräuer H, Forster D, Dietz E, Daschner F, Rüden H. A quality management project in 8 selected hospitals to reduce nosocomial infections: A prospective, controlled study. Infection Control and Hospital Epidemiology. 2002;23:91–7.

6. European Centre for Disease Prevention and Control. European surveillance of healthcare-associated infections in intensive care units - hai-net icu protocol, version 1.02. Stockholm: ECDC; 2015.

7. “European Centre for Disease Prevention and Control”. “ECDC annual epidemiological report for 2015”. “ECDC Stockholm”; 2017. Available from: https://ecdc.europa.eu/sites/portal/files/documents/AER_for_2015-healthcare-associated-infections_0.pdf

